# The genetics of resistance to *Morinda* fruit toxin during the postembryonic stages in *Drosophila sechellia*

**DOI:** 10.1101/014027

**Authors:** Yan Huang, Deniz Erezyilmaz

## Abstract

Many phytophagous insect species are ecologic specialists that have adapted to utilize a single host plant. *Drosophila sechellia* is a specialist that utilizes the ripe fruit of *Morinda citrifolia*, which is toxic to its sibling species, *D. simulans*. Here we apply multiplexed shotgun genotyping and QTL analysis to examine the genetic basis of resistance to *M. citrifolia* fruit toxin in interspecific hybrids. We find that at least four dominant and four recessive loci interact additively to confer resistance to the *M. citrifolia* fruit toxin. These QTL include a dominant locus of large effect on the third chromosome (QTL-III_*sim*_a) that was not detected in previous analyses. The small-effect loci that we identify overlap with regions that were identified in selection experiments with *D. simulans* on octanoic acid and in QTL analyses of adult resistance to octanoic acid. Our high-resolution analysis sheds new light upon the complexity of *M. citrifolia* resistance, and suggests that partial resistance to lower levels of *M. citrifolia* toxin could be passed through introgression from D. sechellia to *D. simulans* in nature. The identification of a locus of major effect, QTL-III_*sim*_a, is an important step towards identifying the molecular basis of host plant specialization by *D. sechellia*.

## Introduction

How genetic architecture may contribute to the emergence of ecological specialization remains an open question (Forister et al. 2012). Genetic mapping methods between recently diverged species have made it possible to assess the relative contributions of dominance, epistasis, complexity and effect size upon the emergence of adaptive traits. The fruit fly, *Drosophila sechellia* provides a clear example of adaptive specialization within the *Drosophila simulans* clade, which includes D. simulans, D. sechellia and D. mauritiana. The three species emerged on islands in the Indian Ocean. *D. simulans* probably evolved on Madagascar, but *D. sechellia* and *D. mauritiana* are endemic to the Seychelles and Mascarene Islands, respectively (Lachaise et al. 1986). Molecular analysis suggests that *D. simulans*, *D. mauritiana* and *D. sechellia* diverged simultaneously about 242,000 years ago, although subsequent admixture has occurred since speciation (Kliman et al. 2000; Garrigan et al. 2012). While both *D. simulans* and *D. mauritiana* exploit a wide variety of fruit, *D. sechellia* is a specialist, and wild *D. sechellia* are found, most frequently, on the fruit of *Morinda citrifolia* (Tsacas and Bachli, 1986; Louis and David, 1986; Matute and Ayroles, 2014), which is toxic to other species of *Drosophila* when it is ripe (Louis and David 1986; R’Kha et al. 1991; Legal et al. 1994). Adult *D. sechellia* are attracted to ripe *M. citrifolia*, and oviposition behavior is stimulated by *M. citrifolia* fruit volatiles (R’Kha et al. 1991; Legal et al. 1992; Higa and Fuyama. 1993; Legal et al. 1999). Neither unripe nor fermenting *M. citrifolia* fruit is toxic to *D. simulans* or *D. melanogaster* (Legal et al. 1992; Farine et al. 1996; Legal et al. 1994). Gas chromatography-mass spectrometry analysis of ripe *M. citrifolia* fruit revealed an abundance of the compound, octanoic acid, a linear 8-chain fatty acid that was prevalent in ripe, but not unripe or fermenting fruit (Legal et al. 1994; Farine et al. 1996; Legal et al. 1999). Octanoic acid is toxic to *D. simulans*, *D. mauritiana* and *D. melanogaster* at all stages of development (R’Kha et al. 1991; Legal et al. 1992; Jones 1998; 2001).

The genetic basis for octanoic acid resistance varies with developmental stage. Using 15 genetic markers, Jones (1998) found that resistance in adults is mostly dominant, and is conveyed by a region on 3R, two regions on the X and additional factors on the second chromosome that could not be localized. To identify larval resistance genes, Jones (2001) used eleven genetic markers and found a dominant major effect locus on the right arm of the third chromosome, with smaller effect loci on 2L and 2R, as well as sex-specific epistatic interactions. A scan for recessive loci revealed a single locus of large effect on 3R and a small-effect interaction between 2L and 2R. Although these analyses have established that octanoic acid resistance in *D. sechellia* is a trait of moderate complexity, the low resolution of these analyses makes it difficult to guess the number or location of loci that are involved. It is therefore not possible to compare the recessive and dominant factors, or adult and juvenile resistance loci with confidence. In addition, mapping at lower resolution increases the likelihood of finding ‘ghost QTLs’; QTL peaks created between two true loci when intervening markers are not available to resolve them (Broman and Speed, 1999). With the emergence of low cost next generation sequencing, it has become possible to assign ancestry at each polymorphic site that exists between two genomes. DNA barcoding methods make high-density genotyping cost effective by multiplexing many individual genomes into a single sequencing library (Baird et al. 2008; Andolfatto et al. 2011). In order to create a high resolution picture of the genetic architecture of *M. citrifolia* resistance in *D. sechellia*, we have used multiplexed shotgun genotyping (MSG), to genotype *M. citrifolia* resistant and sensitive larvae at hundreds of thousands of markers. Here we show that the genetic basis for larval resistance in *D. sechellia*, from the second larval instar (L2) to puparium formation, is conferred through a single dominant locus of large effect in combination with smaller-effect dominant and recessive loci.

## Materials and Methods

### Larval Resistance Assays

To generate synchronized larvae, we transferred individual late first instar (L1) larvae from timed egg lays to separate media dishes. After 2 hours, L2 larvae were identified and then moved to *M. citrifolia*, the octanoic acid test medium, or to regular fly food. The midpoint of the 2-hour interval is taken as the time of L1-L2 molt. Molting larvae were identified by the presence of double mouth hooks, posterior to anterior peristalsis movements or ecdysis.

To generate the octanoic acid test medium, octanoic acid from Sigma, (St. Louis, MO) was diluted in water and mixed with 0.25 g nutritional yeast and 2.0g of Carolina instant *Drosophila* food, formula 4-24 (Burlington, NC), and used within one hour. For the octanoic acid dose-response tests, larvae were staged at the L1 to L2 molt, and moved to octanoic acid medium three hours after the onset of the second instar to avoid handling during the L2 molt. The number of pupae was recorded after three days. Larvae were tested in groups of 30.

*M. citrifolia* was grown in Hawaii, and shipped frozen overnight to New York (a generous gift of Scot Nelson, University of Hawaii). We find that each fruit from this source always produces 100% lethality in *D. simulans*. For the *M. citrifolia* resistance assays used to phenotype larvae for QTL mapping, L2 larvae were individually selected from 6-12 hour collections and transferred to thawed *M. citrifolia* fruit that had been frozen at the ‘transluscent gray’ stage while still firm (Chan-Blanco et al. 2006). Frozen fruits were thawed overnight at room temperature and used within 24 hours. The larvae on *M. citrifolia* were monitored for the next 6 hours. Those larvae that stopped moving, (but were not undergoing peristalsis movements of a molt) were considered *M. citrifolia* sensitive. Those that survived to form puparia were considered resistant.

### Genotyping

We generated recombinant mapping populations by crossing female *D. simulans^Nueva^* to male *D. sechellia^w30^*, since the reciprocal cross does not produce offspring in our experience. The virgin F1 hybrid females were backcrossed to either parental stock.

For multiplexed shotgun genotyping, DNA was extracted from individual flies using a modified version of the Purgene protocol (supporting information). Custom barcode adapters (supporting information) were ligated onto genomic DNA, and libraries of 384 barcoded individual genomes were processed according to Andolfatto et al. (2011). For our *D. simulans* backcross, we combined barcoded genomic DNA from 384 individual sensitive larvae into a lane of an Illumina HiSeq (SRS645327), and 384 resistant larvae into another lane of an Illumina HiSeq (SRS645328). For the *D. sechellia^w30^* backcross, we combined barcoded DNA from 186 resistant larvae with 192 sensitive larvae into a separate lane (SRS645326). The reads of all our libraries were mapped to a recent *D. simulans* genome assembly (Hu et al. 2012) that was updated with the parental strains, *D. simulans^Nueva^*(SRS643607) and *D. sechellia^w30^* (ID: 2867884) with the UpdateParental function of the MSG package (Github:JaneliaSciComp/msg). The reference genome (Hu et al. 2012) uses the *D. melanogaster* genome release r5.33 as a reference guide, and we use these *D. melanogaster* coordinates throughout the manuscript. The MSG package was installed on Galaxy Cloudman on the Amazon Elastic Compute Cloud. Configuration files, msg.cfg and update.cfg with selected parameters are available as supporting information.

We used indel locations reported in Erezyilmaz and Stern (2013) to identify species-specific insertions and deletions (supporting information). The primers 3R22.36f: 5’ CAGTACACAATGGTGGGCAT 3’, and 3R22.36r: 5’TTTGGTCCAAAAGGAAGCTGA 3’ straddle a region of a *D. simulans* - *D. sechellia* alignment that contains a 20bp deletion in *D. simulans*. The PCR-amplified products were visualized on a gel made with 2.5% Lonza MetaPhor agarose (Basel, Switzerland).

### Post MSG processing

The perl script, pull_thin.pl created by David Stern (Janelia Farm and HHMI; supporting information) was used to prune our genotype files to relevant locations that straddle recombination events. The *D. sechellia^w30^* backcross data set was thinned from 406,475 markers to 8,206 markers. The *D. simulans* backcross dataset was thinned from 298,768 markers to 7,483 markers.

### Qtl Mapping

We treated *M. citrifolia* resistance as a binary trait and scored survivorship to puparium formation. We used the *detectable* and *powercalc* functions of the R package, R/QTL Design (Sen et al. 2007) to determine the power of our QTL experiments to detect given effect sizes. We used the *scanone* function in R/qtl (Broman et al. 2003) to scan for QTL, and plotted the results in R. To assess the statistical significance of our LOD scores in the one-dimensional analyses, we performed permutation analysis with 1000 replicates. For two-dimensional scans, we used reduced datasets that contained 190 markers for the *D. sechellia^w30^* backcross, and 125 markers for the *D. simulans* backcross. We analyzed the reduced datasets using the *scantwo* function in R/qtl and in the R/qtl interface, J/qtl (Jackson Labs, Maine). The statistical significance of our two-dimensional scans was assessed using 500 permutation replicates. The fit of models were tested using the *fitqtl* function of R/qtl (Broman et al. 2003).

### Fly Strains

The QTL experiments were performed with a white eyed mutant of the srain, *D. simulans^Nueva^* (San Diego Stock Center # = 14021-0251.006) and *D. sechellia^w30^* (14021-0248.30). The mutant strains, *D. simulans^cutsy,ro,ca^* (San Diego Stock Center # = 14021-0251.116), and *D. simulans^jv,st,e,p^* (San Diego Stock Center # = 14021-0251.174) were used in marker association experiments. We used an inbred line of *D. sechellia* (San Diego Stock Center # = 14021-0248.13), *D. sechellia^D1Alc^* for the marker association experiments with *D. simulans^cutsy,ro,ca^*.

## Results

### Resistance during the late larval stages and puparium formation

Ripening in *M. citrifolia* occurs as hard green fruit turns pale yellow, softens, and becomes a translucent gray color as the fruit ripens (Chan-Blanco et al. 2006). After hatching, *Drosophila* progress through three larval stages (L1-L3). The onset of metamorphosis occurs as the third instar larva becomes immobile and forms a puparium. We used fruits at the ‘translucent gray’ stage, and devised a high-throughput method for selecting juvenile *Drosophila* for resistance. Staged L2 larvae are transferred from fly food to *M. citrifolia* fruit, and allowed to feed and form puparia. Under these conditions, 73% of *D. sechellia^w30^* larvae (N= 180), but no *D. simulans^Nueva^* larvae (N=240), survived to form puparia (Figure 1).

**Figure 1.**
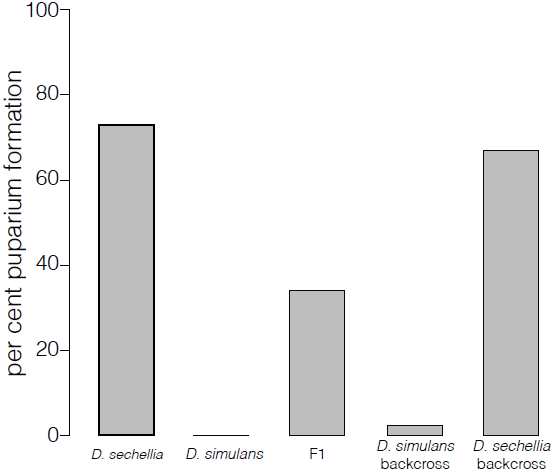
Survival to puparium formation of larvae transferred to ripe *M. citrifolia* fruit. F1 = *D. simulans*^Nueva^/*D. sechellia^w30^* hybrids. *D. simulans* backcross = progeny from F1 females and to *D. simulans*^Nueva^ males. *D. sechellia* backcross = progeny from F1 females and *D. sechellia^w30^* males.

To compare the efficacy of our *M. citrifolia* fruit assay to the effects of pure octanoic acid, we examined the dose-response relationship between octanoic acid concentrations in food for the stages from L2 to pupariation. *D. sechellia^w30^* had a ~ 3-fold greater resistance to octanoic acid than *D. simulans^Nueva^* (Figure 2; D. *sechellia^w30^* LD_50_ = ~0.6% for, *D. simulans^Nueva^* LD_50_ = ~0.22% octanoic acid). The level of toxicity at which 73% of *D. sechellia^w30^* larvae died, roughly corresponds to 0.46% octanoic acid.

**Figure 2.**
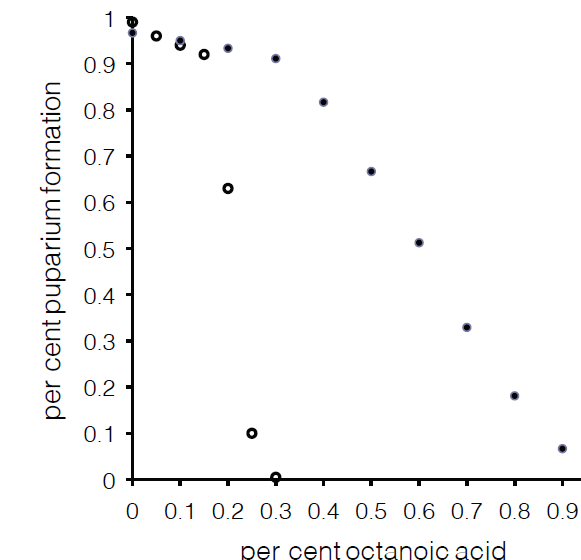
Percentage of larvae that form puparia after transfer to octanoic acid during the second larval instar. The percentage of octanoic acid within *Drosophila* food is given on the X-axis. Filled circles represent 2,070 staged *D. sechellia^w30^* larvae were tested in groups of 30. Between 60 and 240 larvae were used for each data point. Open circles indicate data for 1,620 *D. simulans*^Nueva^ larvae. Between 180 and 270 larvae were used for each data point.

Most L2 larvae did not die upon contact with octanoic acid. Lethality, instead, occurred at a range of times after contact. We tested, next, the possibility that toxicity in *D. simulans* larvae is due to octanoic acid during specific developmental intervals. We exposed *D. simulans^Nueva^* staged L2 larvae to an intermediate dose 0.2 % of octanoic acid for 6 hour periods, then returned larvae to octanoic acid-free medium and measured survivorship to the pupal stage (Figure 3). We found that the survivorship of *D. simulans^Nueva^* larvae during the intermolt period (hours 6-24) was comparable to the survivorship of control larvae that were similarly handled, but not exposed to octanoic acid (Figure 3). In contrast, all *D. simulans^Nueva^* larvae that were exposed to octanoic acid during either the L1 to L2 molt (N = 90), or the L2 to L3 molt (N = 90), did not survive to form puparia (Figure 3). These data show that, at the concentration that we describe here, octanoic acid toxicity during the second larval instar occurs at molts.

**Figure 3.**
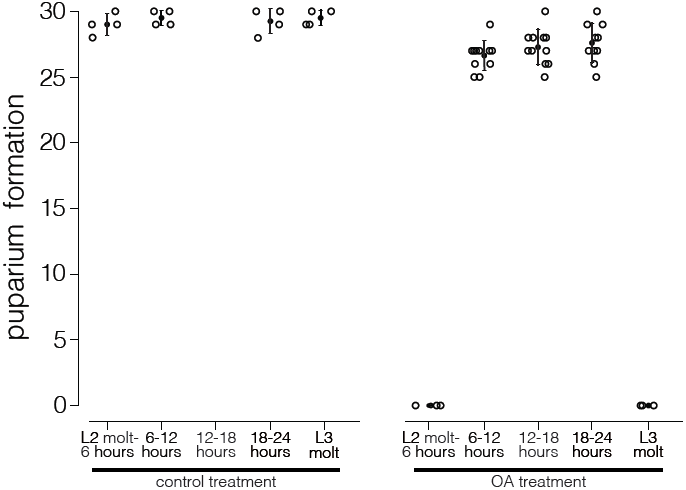
Survivorship to puparium formation after 6 hour treatments during the second larval instar for control (left panel) and 0.2% octanoic acid (OA) treatments. Each open circle represents a test group of 30 larvae. Filled circle indicates the mean and the error bars show standard errors.

### QTL Analysis

D. sechellia and *D. simulans* produce fertile f emale offspring when crossed (Lachaise et al. 1986). Previous work has shown that resistance to *M. citrifolia* toxin by *D. sechellia* adults is mostly dominant in F1 hybrid *D. simulans*/*D. sechellia* larvae, but that resistance to *M. citrifolia* toxin and octanoic acid in developing embryos involves both dominant and recessive factors, as well as a maternal contribution (R’Kha et al. 1991; Jones, 2001). We asked whether resistance is dominant or recessive in our assay. The overall survivorship of F1 *D. sechellia^w30^*/ *D. simulans^Nueva^* hybrids to *M. citrifolia* toxin is intermediate between the survivorship of the two parents, which suggests that resistance in *D. sechellia* is comprised of both dominant and recessive factors (Figure 1). To map these loci at high resolution, we screened L2 larvae from backcrosses to either parent in our *M. citrifolia* assays, and genotyped individual larvae using multiplexed shotgun genotyping (Andolfatto et al. 2011).

We first examined the contribution of recessive loci to *M. citrifolia* resistance in 359 individuals from an F1 backcross to *D. sechellia^w30^*. We find that the map of recessive loci for *M. citrifolia* resistance is dominated by QTL on the third chromosome; all regions of the third chromosome are significant at the 99% level (Figure 4). The highest peak is on the left arm of chromosome III at ~15,860,000 (LOD=20.2). Inclusion of a neighboring peak at ~3:18,900,000 did not increase the likelihood of our model significantly (Table 1). We therefore treat this region as one locus, QTL-III_*sec*_a. A second large effect locus appears on the right arm at ~3:40,070,000 with LOD =17.0 (QTL-III_*sec*_b, Figure 4). In addition, 2 QTL scans indicate that a resistance locus exists on chromosome III at ~45,180,000 (QTL-III_*sec*_c, LOD = 10.9; Figure 4, S1), and inclusion of this region improves the fit or our model (Table 1). We also found a significant contribution from the X chromosome; the left half is significant at the 95% confidence level, and a peak lies at ~X:10,680,000 (QTL-X_*sec*_, LOD = 6.5). Recessive loci on the second and the fourth chromosomes did not contribute significantly to resistance. The best-fit model contains QTL-III_*sec*_a, QTL-III_*sec*_b, QTL-III_*sec*_C and QTL-X_*sec*_ interacting additively (Table 1). We did not detect significant epistasis between loci in our two-QTL scans (Figure SI).

**Figure 4.**
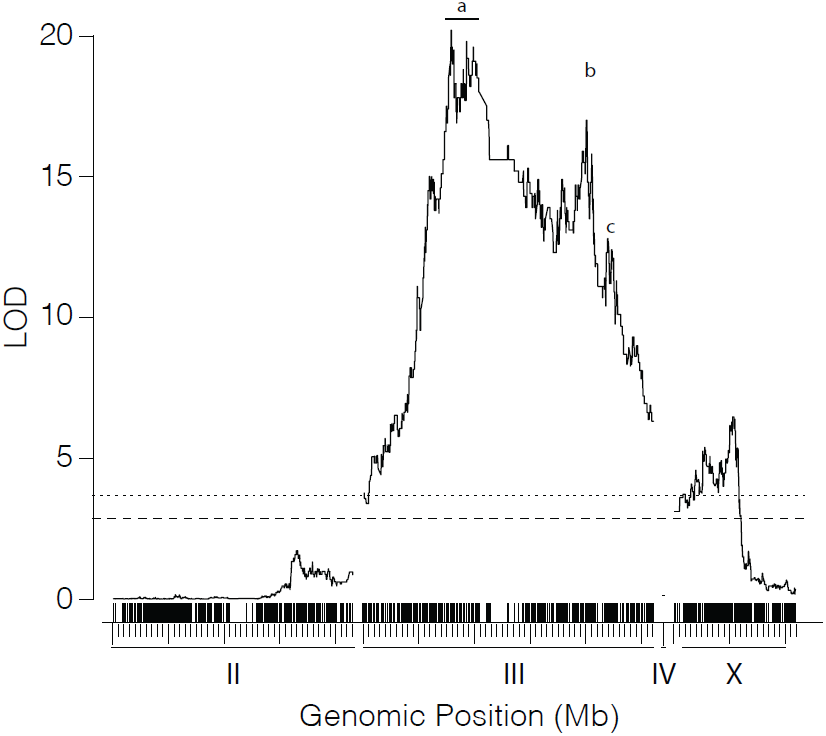
QTL map of the recessive loci for *M. citrifolia* resistance in the*D.sechellia^w30^* backcross. The log of the likelihood ratio (y axis) is plotted against physical distances (x axis). The locations of markers is shown in the upper row of vertical ticks. The genomic locations are given in the bottom row ofvertical ticks; each large tick is 10Mbp, and the smaller ticks show 1 Mbp intervals. Dashed, and 28 dotted lines indicates the threshold for LOD scores at the 0.05 and 0.01 significance levels, respectively. The locations of QTL-III_*sec*_a (a), QTL-III_*sec*_b (b), and QTL-III_*sec*_c (c) are indicated.

**Table 1.** QTL models for the *D. simulans* and *D. sechellia* backcrosses.

| Cross | Number of QTLs in Model | LOD of model | Percent of Variance Explained | QTL locations | Percent of species difference | LOD drop one<sup>1</sup> |
| --- | --- | --- | --- | --- | --- | --- |
| *D. sechellia* backcross | 2 | 27.9 | 41.2 | 3 ~ 15.9 Mbp<br>3 ~ 39.0 Mbp | 15.2<br>10 | 11.5<br>7.3 |
| *D. sechellia* backcross | 3 | 31 | 45.2 | X ~ 10.8 Mbp<br>3 ~ 15.9 Mbp<br>3 ~ 39.0 Mbp | 5.9<br>12.3<br>9.2 | 4.9<br>9.7<br>7.3 |
| *D. sechellia* backcross | 4 | 31.4 | 45.5 | X ~ 10.8 Mbp<br>3 ~ 15.9 Mbp<br>3 ~ 39.0 Mbp<br>3 ~ 19.0 Mbp | 6.2<br>1.7<br>7.5<br>0.6 | 5.1<br>1.4<br>6.2<br>0.5 |
| *D. sechellia* backcross | 4 | 33.8 | 48.2 | X ~ 10.8 Mbp<br>3 ~ 15.9 Mbp<br>3 ~ 39.0 Mbp<br>3 ~ 45.2 Mbp | 5.7<br>13.4<br>2.0<br>3.4 | 5.0<br>11.0<br>1.7<br>2.9 |
| *D. simulans* backcross | 2 | 37 | 28.4 | X ~14.4 Mbp<br>3 ~47.0 Mbp | 3.0<br>24.0 | 4.4<br>31.7 |
| *D. simulans* backcross | 2 | 38 | 28.7 | 2 ~ 43.5 Mbp<br>3 ~ 47.0 Mbp | 3.6<br>24.1 | 5.2<br>32.4 |
| *D. simulans* backcross | 3 | 41.9 | 31.4 | X ~ 14.4 Mbp<br>2 ~ 43.5 Mbp<br>3 ~ 47.0 Mbp | 2.7<br>3.2<br>22.7 | 4.1<br>4.7<br>31.4 |
| *D. simulans* backcross | 4 | 45.8 | 33.9 | X ~14.4 Mbp<br>2 ~ 43.5 Mbp<br>3 ~ 47.0 Mbp<br>3 ~ 9.0 Mbp | 2.6<br>3.1<br>21.8<br>2.5 | 4.1<br>4.8<br>30.9<br>3.8 |
| *D. simulans* backcross | 5 | 46.3 | 34.2 | X ~14.4 Mbp<br>2 ~ 43.5 Mbp<br>3 ~ 47.0 Mbp<br>3 ~ 9.0 Mbp<br>3 ~ 5.0 Mbp | 2.7<br>3.0<br>21.8<br>0.3<br>0.4 | 4.2<br>4.6<br>30.9<br>0.5<br>0.6 |

<sup>1</sup> Log likelihood ratios comparing the full model to a model with the specified QTL removed.

Given the genetic and environmental variances of our lines, and the number of backcross individuals tested, our *D. sechellia^w30^* backcross QTL experiment is designed to detect effect sizes of 0.19 and 0.175 with 90, and 80 per cent power, respectively. The effect sizes of QTL-III_*sec*_a, QTL-III_*sec*_b, QTL-III_*sec*_c and QTL-X_*sec*_ exceed the threshold for detection with 90% power (Figure S2).

We next examined the effect of dominant loci in D. sechellia’s resistance to *M. citrifolia* toxin in progeny of F1 backcrosses to *D. simulans^Nueva^*. QTL analysis showed that resistance to *M. citrifolia* is dominated by a QTL of large effect on the right arm of chromosome III at ~ 46,954,562 bp (QTL-IIIa_*sim*_; LOD = 33.8; Figure 5). Larvae with one copy of the *D. sechellia^w30^* allele at QTL-III_*sim*_a were more than twice as likely to survive to pupariate (Figure 5B; Figure S2). We also detected two significant peaks on the left arm of chromosome 3 at ~ 4,898,627 bp (LOD=4.42) and at 9,012,208 bp (LOD = 4.48), although a model including both markers did not significantly increase the likelihood score over a model that includes just one of the two 3L markers. We therefore treat these terms as a single locus, QTL-III_*sim*_b. In addition, a broad region of significance on the right arm of chromosome II has a peak at ~43,480,964 bp (QTL-II_*sim*_, LOD = 5.8) and spans several megabases. Finally, we found a broad region on the X chromosome between ~X:10,000,000 and ~X: 18,200,000 that has a peak at ~X: 14,358,268 bp (QTL-X_*sim*_, LOD= 6.1). We did not detect significant epistasis between QTLs in two-dimensional, two QTL scans (Figure S1). Instead, we find that a model that fits our data best consists of QTL-IIIa_*sim*_, QTL-IIIb_*sim*_, QTL-II_*sim*_ and QTL-X_*sim*_ interacting additively (Table 1). Inclusion of additional QTL did not improve the fit of our model significantly (Table 1).

**Figure 5.**
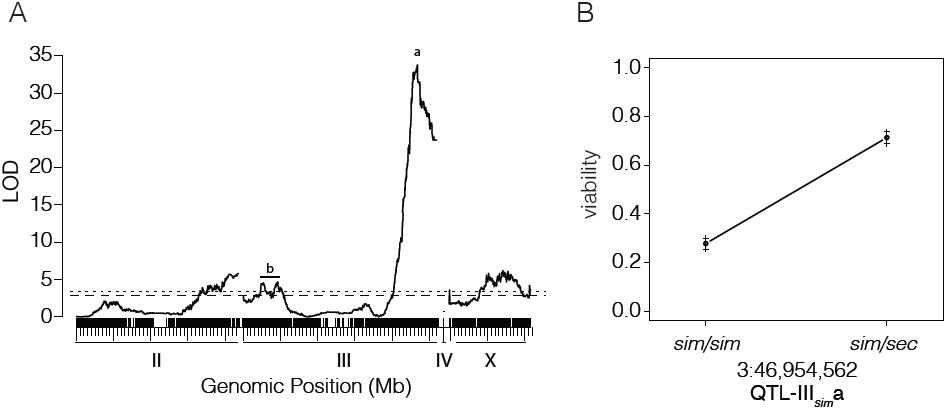
QTL map of the dominant loci for *M. citrifolia* resistance in the *D.simulans*^Nueva^ backcross. (A) LOD scores vs genomic position for the four chromosomes. The upper rug of ticks shows the location of markers, while the lower tick marks indicate 1 and 10 Mbp intervals along each chromosome. The significance thresholds for the 0.05 and 0.01 levels of the LOD scores are shown by the dashed and dotted lines, respectively. The location of QTL-III_*sim*_a (a) and QTL_III_*sim*_b (b) are indicated above the QTL peaks on chromosome III. (B) Effect plot for QTL-III_*sim*_a showing the phenotypic effect of substituting a *D. simulans*^Nueva^ allele (*sim*) for a *D. sechellia^w30^* allele (*sec*) at the marker 3:46,954,562, where 0 = lethality, and 1 = survivorship to puparium formation.

Based upon the sample size, environmental and genetic variances of our cross, a QTL with effect size of 0.12, where 0 is sensitive and 1 is resistant, would be detectable with 80% power, and a QTL with effect size of 0.134 would be detected with 90% power in our *D. simulans^Nueva^* backcross. The effect sizes of the four QTL that we report exceed the 90% power threshold (Figure S2).

### Tests of QTL-III_*sim*_a and the effect of the X chromosome

The location of our largest-effect QTL in the *D. simulans* ^*Nueva*^ backcross differs from the large effect locus described by Jones for octanoic acid sensitivity in larvae (2001) and in adults (1998). We therefore performed additional checks of our approach. First, we compared the genotype of a sample of our MSG-genotyped flies at ~3:46,689,564 with an indel marker at 3:46,617,828 using PCR. The PCR-indel genotypes of all 179 genotyped individuals matched the genotypes produced by MSG (data not shown). To confirm the existence of QTL-III_*sim*_a, we next crossed a *D. simulans* strain, marked by the *cutsy* mutation and by claret (ca) at ~3:50,000,000, (Kimble and Church, 1983) to *D. sechellia^D1A1C^*, and backcrossed F1 females to the marked *D. simulans^cutsy,ca^* parental strain. The genetic map position of *cutsy* is 3-74 in *D. simulans* (Coyne, 1997) which is, roughly, ~3:42,000,000 (*D. melanogaster* genome location). We compared the frequency of each recessive marker in 242 F1 backcross progeny that were viable in *M. citrifolia* from L2 to adulthood with the frequency of each marker in 872 larvae that were not exposed to *M. citrifolia* fruit. We found that *cutsy^+^* (∑^2^ = 20.6, *p* = 5.8 × 10^−6^), was most strongly associated with *M. citrifolia* resistance, followed by *ca^+^* (∑^2^ =6.2, *p* = 0.01). We next used the molecular markers at 3:46,617,828 to test for linkage to an additional 89 resistant and 91 sensitive *D. simulans* backcross progeny. We found strong linkage to this region (*p* = 4 × 10-5, Fisher’s Exact Test; ∑^2^ = 27, *p* = 2.06 × 10^−7^, Chi-squared test, Table 2).

**Table 2.** linkage of markers on 3R to *M. citrifolia* resistance.

| location<sup>i</sup> | marker<sup>ii</sup> | observed | expected | $\Sigma^2$ | $p$ – value<sup>iii</sup> |
| --- | --- | --- | --- | --- | --- |
| ~3:42,000,000 | *cutsy*<sup>+</sup> | 156 | 121 | 20.6 | $5.8 \times 10^{-6}$ |
|  | *cutsy*<sup>-</sup> | 86 | 121 |  |  |
| 3:46,617,828 | *indel*<sup>e</sup> | 69 | 46 | 27.0 | $2.1 \times 10^{-7}$ |
|  | *indel*<sup>i</sup> | 20 | 44 |  |  |
| ~3:50,000,000 | *ca*<sup>+</sup> | 137 | 118 | 6.2 | 0.01 |
|  | *ca*<sup>-</sup> | 105 | 124 |  |  |

<sup>i</sup>Location in the *D. melanogaster* reference genome.
<sup>ii</sup>For *cutsy* and *claret* (*ca*), inheritance of a second copy of the recessive marker from *D. simulans*. *Indel*<sup>e</sup> indicates inheritance of the *D. sechellia* variant of the indel polymorphism, *indel*<sup>i</sup> indicates the inheritance of a second copy of the *D. simulans* indel polymorphism.
<sup>iii</sup>Chi-squared test.

These data confirm that QTL-III_*sim*_a, is located near ~3:47,000,000, and suggest that QTL-III_*sim*_a may be closer to ~3:42,300,000 than to ~3:50,000,000 (Table 2). Finally, we asked if *M. citrifolia* resistance in our assay is linked to *ebony* (*e*), since Jones et al (2001) found that a QTL linked to e confers resistance to octanoic acid in a larval assay. The genomes of the *D. simulans* clade share a large inversion on 3R with respect to *D. melanogaster* that contains *e* (Ashburner, 1989). In our QTL map, which uses the D. melanogaster genome as a scaffold, the e gene is located at ~3:41,620,000, but in *D. simulans*, e is located at ~3:28,656,573. Like Jones (2001), we also find that *M. citrifolia* resistance is linked to e in our assay, although the linkage is weaker than the linkage between resistance and *cutsy*, *ca or* 3:46,617,828 ( Table 3; ∑^2^ = 5.08, *p* = 0.02).

**Table 3.**
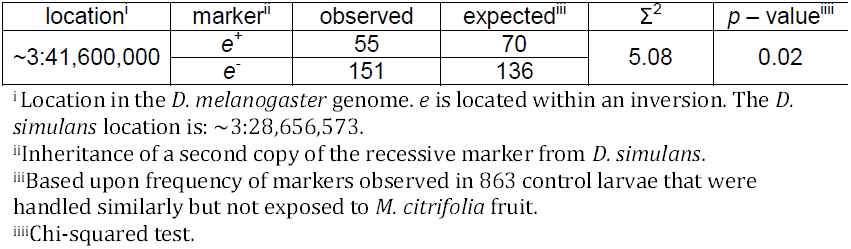
*M. citrifolia* linkage to *ebony* (*e*).

| location<sup>i</sup> | marker<sup>ii</sup> | observed | expected<sup>iii</sup> | $\Sigma^2$ | *p* – value<sup>iiii</sup> |
| --- | --- | --- | --- | --- | --- |
| ~3:41,600,000 | *e*<sup>+</sup> | 55 | 70 | 5.08 | 0.02 |
|  | *e*<sup>-</sup> | 151 | 136 |  |  |

<sup>i</sup> Location in the *D. melanogaster* genome. *e* is located within an inversion. The *D. simulans* location is: ~3:28,656,573.
<sup>ii</sup> Inheritance of a second copy of the recessive marker from *D. simulans*.
<sup>iii</sup> Based upon frequency of markers observed in 863 control larvae that were handled similarly but not exposed to *M. citrifolia* fruit.
<sup>iiii</sup> Chi-squared test.

Our analysis of both the *D. simulans* backcross and the *D. sechellia^w30^* backcross revealed resistance loci on the X chromosome, which contradicts previous work (Jones, 2001). If the X chromosome does confer resistance in our assay, larvae with one X chromosome from *D. sechellia^w30^* should have greater resistance than larvae with only a *D. simulans* chromosome. As an additional test of our QTL model, we asked if female F1 larvae had greater survivorship than male F1 larvae from the cross, ♀ *D. simulans^Nueva^* × ♂ *D. sechellia^w30^*, since males would only have a single X chromosome from *D. simulans^Nueva^* mother. In this case we find that more F1 females than F1 males (*p* = 9.36 × 10^−7^, Fisher’s Exact Test, N = 141 larvae in *M. citrifolia* vs N = 371 control larvae). This result indicates that the X chromosome from *D. sechellia* has a significant effect upon resistance to *M. citrifolia* toxin.

## Discussion

We have addressed the genetic basis for evolved resistance to *M. citrifolia* fruit toxin in *D. sechellia*. We find that resistance to the toxic fruit during the period from the second instar to puparium formation is composed of both dominant and recessive effects. Multiple significant recessive loci were detected on the third chromosome (QTL-III_*sec*_a-c), as well as a region on the X chromosome (QTL-X_*sim*_). A scan for dominant loci identified a large effect locus at ~3: 46,954,562 bp (QTL-IIL_*sim*_a) and three regions of smaller effect that are located on the X (QTL-X_*sim*_), on 2R(QTL-II_*sim*_), and on 3L (QTL-IIL_*sim*_b). QTL-IIL_*sim*_ a is within an interval that overlaps with QTL-III_*sec*_C, which may indicate that the two QTL are the same locus that acts additively. We found no evidence for significant epistasis between any QTLs. Additional checks of our locus of largest effect, QTL-III_*sim*_a using visible and molecular markers confirms that the locus that confers resistance to *D. sechellia^w30^* is between ~3:42,000,000 and ~3:47,000,000, but closer to ~3:47,000,000.

Although the set of loci that we detected overlap with the regions identified by Jones (2001), the two sets are not identical. Both analyses indicate a major effect locus on 3R, (Figure 5, Table 1), but Jones (2001) finds that the major effect locus is near e, and a region near QTL-III_*sim*_a was not significant. Another major difference between our QTL map and that of Jones (2001) is the effect of the X chromosome. We found significant QTL peaks on the X chromosome in both the *D. simulans* and *D. sechellia^w30^* backcrosses, while Jones (2001) did not detect any effects on the X chromosome. In addition, Jones found that a region on the left arm of chromosome II accounted for 11-12% of the phenotypic variation in the *D. simulans* backcross (Jones, 2001), but we did not detect significant QTL on 2L in either backcross (Figure 4, 5). Finally, Jones (2001) detects epistasis between 2L and 2R in the *D*. *sechellia* backcross and between 2R and 3R in the *D. simulans* backcross whereas we did not detect significant epistasis between loci.

Several factors may account for the discrepancies between our QTL map and the previous analysis by Jones (2001). Firstly, we used fresh-frozen *M. citrifolia* for our assays, while Jones used purified octanoic acid in fly food. Although previous work (Legal et al, 1994) has established that octanoic acid is the toxic component of *M. citrifolia* fruit, other components of the fruit may contribute to the toxicity or uptake of octanoic acid. Secondly, the two analyses may test different phases of development. Jones (2001) allowed females to oviposit on control or octanoic acid containing medium, then recorded the visible markers of emerging adults. *D. simulans* embryos are highly sensitive to *M. citrifolia* (R’Kha, 1991), while *D. sechellia* embryos are largely resistant. The lethality of octanoic acid decreases over time, presumably as the semi-volatile compound dissipates. In Jones’ assay, therefore, embryonic resistance factors would be under the greatest selection while later stages of development would experience increasingly lower effective octanoic acid concentrations. Jones (2001) also provides strong evidence for a maternally inherited resistance factor, which further complicates genetic mapping. A locus that encodes a maternally-provided factor would not appear in selection experiments if the offspring have resistant mothers. Therefore, any differences between the set of larval resistance factors described here and those described by Jones (2001) could be due to maternally provided factors, selection upon different stages of resistance, or any combination of the two. Finally, we compare the genotypes of resistant larvae with those of sensitive larvae for our QTL analyses, whereas Jones compares the genotypes of resistant larvae with those of larvae that have not been exposed to octanoic acid. Interestingly, linkage to the region containing e was not significant in our QTL map, but it was significant when we used untreated backcross progeny as a control (Figure 5; Table 3).

In contrast to larval resistance, the resistance to volatile octanoic acid in adults is conferred through dominant loci on the 2^nd^, 3^rd^ and X chromosomes (Jones, 1998). The resistance factor on chromosome 2 had too small of an effect to be mapped, but the loci on the X and III were further resolved with visible markers (Jones, 1998). A region on the right arm of chromosome 3 that is linked to *e* had the greatest effect, and recent work has fine mapped this locus to the interval bounded by ~3:26,136,000 and ~3:26,315,000 (Hungate et al. 2014). Interestingly, our *D. simulans* backcross QTL map for larval resistance overlaps with a locus on the X chromosome discovered by Jones (1998) for adult resistance; a region between miniature (~X:11,700,000) and forked (~X:17,130,000). We find a broad significant region from ~X:10,000,000 to ~X:18,200,000 that peaks at ~X:14,358,268.

The response of insect populations to insecticide treatment may provide insight into how resistance to *M. citrifolia* toxin may have evolved in *D. sechellia*. Exposures to insecticide concentrations that lie within the distribution of viability tend to produce resistance that is based on multiple loci, each of small effect (McKenzie et al, 1992). Such variation has been found to regulate the expression level or copy number of detoxifying enzymes, such as the cytochrome P450s and by other metabolic enzymes, such as carboxylases and esterases (McKenzie and Batterham, 1994; Ranson et al. 2002; ffrench-Constant et al. 2004). Selection outside of the viability distribution with very high levels of insecticide, on the other hand, tends to produce large-effect loci conferred by amino acid substitutions of single genes that are the targets of pesticides. These targets include ligand-gated ion channels, like the GABA receptor subunit in which an amino acid replacement confers resistance to dieldrin (ffrench-Constant et al. 1991), or a voltage gated sodium channel in which resistance to DDT is conferred by either of two amino acid replacements (Williamson et al. 1996; Miyazaki et al. 1996). The genetic architecture of resistance to *M. citrifolia* fruit toxin in *D. sechellia* that we describe here bears hallmarks of both types of selection; one large effect locus on 3R accounts for 24% of the phenotypic difference between *D. simulans^Nueva^* and *D. sechellia^w30^*, but in the best-fit model, additional smaller effect loci also confer resistance (Table 1).

Data from selection experiments suggest that complete resistance did not arise only through consolidation of existing variation in resistance alleles alone. Colson (2004) selected for increased resistance in a cosmopolitan strain of *D. simulans* for 20 generations. She found that resistance to octanoic acid rapidly increased by 18% before it plateaued at a fraction of the resistance seen in *D. sechellia*. These data show that either the variation within the cosmopolitan strain that was used is not representative of the variation within the common *D. simulans* - *D. sechellia* ancestor, or that resistance in *D. sechellia* arose through new mutation(s). Interestingly, two regions discovered in these selection experiments overlap substantially with QTL-II_*sim*_ at cytological location 57C and QTL-X_*sim*_, at cytological location 13D (Colson 2004). Colocalization of our small-effect loci with the evolved resistance regions of Colson (2004) supports a scenario where the smaller-effect loci discovered in our experiments are the products of selection within the viability distribution of D. simualns for octanoic acid.

The genetic complexity of resistance to *M. citrifolia* fruit in *D. sechellia* appears to be a barrier to full introgression of this trait into *D. simulans*. Amlou et al (1997) tried to introgress octanoic acid resistance into *D. simulans* by backcrossing resistant adult hybrids to *D. simulans*. Despite strong directional selection, resistance decreased with each generation to *D. simulans* levels. Our own efforts to introgress the resistance phenotype into *D. simulans^Nueva^* (phenotype-based introgression) did not progress beyond one generation of selection upon ripe *M. citrifolia* (data not shown). However, the lack of epistasis between the four major QTL suggests that partial resistance could easily be introgressed from *D. sechellia* to *D. simulans*. Partial resistance would improve the viability of hybrid flies on fermenting fruit, which has been shown to have lower levels of octanoic acid (Legal et al. 1994; Farine et al. 1996; Legal et al. 1999). Such introgression of partial resistance may occur routinely in the Seychelles, where *D. simulans* and *D. sechellia* co-exist. Matute and Ayroles (2014) recently showed that *D. simulans* and *D. sechellia* hybrids are prevalent in some of the islands in the Seychelles. They also found that F1 *D. simulans*/*D.sechellia* hybrids and morphologically *D. simulans* flies are found on *M. citrifolia* fruit, although the stage of fruit maturation was not reported (Matute and Ayroles, 2014).

QTL analyses typically produce confidence intervals that are too large to resolve candidate genes. In the case of QTL-III_*sim*_a, the confidence interval created by a 1.5 LOD score drop from the peak value of QTL-III_*sim*_a spans from ~3:45,769,000 to ~3:47,000,000, about 1.2 Mbp. This interval contains 128 protein-coding genes, only 70 of which are named. There are also numerous non-protein coding genes, which have recently been shown to regulate an evolved difference in morphology in Drosophila (Arif et al, 2013). Additional work with introgression lines or *D. melanogaster* deficiency strains will be needed to further resolve this interval with fine scale mapping. These experiments are currently underway in our laboratory.

## Acknowledgments

We thank Prof Scot Nelson, (University of Hawaii, Manoa) for his generous gifts of *M. citrifolia* fruit. We also thank Greg Pinero and David Stern for installation of the MSG program onto the Amazon/Galaxy Cloud. This work was supported by Stony Brook University start-up funds.

## Supporting Information

**Figure S1**. Two dimensional, two-QTL scans for the *D. sechellia^w30^* and *D. simulans*^Nueva^ backcrosses. For each graph, the top triangle shows the improvement in fit of the full model over a single QTL model (LOD_fv1_), the bottom right triangle shows the improvement in fit of the additive model (LOD_av1_) over the single QTL model. A) In the *D. sechellia^w30^* backcross, little difference is found between the LOD_fv1_ and LOD_av1_ models. The significant interaction between the two QTL on chromosome 3 is additive: ~3:45,009,000 × ~3:15,860,000 (LOD_fv1_ = 10.0, LOD_av1_ = 9.6, LOD_epistasis_ = 0.38). These locations correspond to QTL-III_*sec*_a and QTL-III_*sec*_c. B) *D. simulans*^Nueva^ backcross. A locus at ~3:46,040,000 interacts with a locus at ~X:15,001,000 (LOD_fv1_ = 7.1, LOD_av1_ = 4.9, LOD_epistasis_ = 2.2). A second interaction occurs between loci at ~2:43,020,000 and ~X:15,001,000 (LOD_fv1_ = 5.7, LOD_av1_ = 4.8, LOD_epistasis_ = 0.9). A third interaction occurs between loci on the right arms of the second and third chromosomes, ~3:47,001,000 × ~2:43,024,000 (LOD_fv1_ = 4.4, LOD_av1_ = 4.4, LOD_epistasis_ = 0.001). A fourth interaction occurs between two loci on the third chromosome, ~3:9,011,000 and ~3:47,001,000 (LOD_fv1_ = 6.1, LOD _av1_ = 3.4, LOD_epistasis_ = 2.7). The significance of all the LOD_fv1_ and LOD_av1_interactions met or exceeded the *p* = 0.01 threshold, but no episatic interactions were significant. The locations of the loci reported here are all within the confidence intervals of QTL-II_*sim*_, QTL-III_*sim*_, QTL-III_*sim*_b and QTL-X_*sim*_

**Figure S2**. Whole genome effect plots for (A) the *D. sechellia^w30^* backcross, (B) the *D. simulans*^Nueva^ backcross. The effect of each location on the autosomes is given in blue, and the standard errors are shaded in light blue. For the X chromosome, effect sizes are given in red, with standard errors shaded in pink. For males, the effect size of each location on the X is given in orange, and the standard errors are shaded in light orange.

**Supporting Information File 1.** The sequence of 384 barcoded adapters that were used to ligate FC2 linkers for Illumina sequencing (Andolfatto et al, 2011).

**Supporting Information File 2.** The configuration file with parameters that was used for updating the *D. simulans^w501^* genome with *D. simulans^Nueva^* and *D. sechellia^w30^* genome reads.

**Supporting Information File 3.** msg.cfg: the configuration file with the parameters that were used for msg mapping.

